# Bactericidal Type IV Secretion System Homeostasis in Xanthomonas citri

**DOI:** 10.1101/647685

**Authors:** William Cenens, Maxuel O. Andrade, Chuck S. Farah

**Affiliations:** Departamento de Bioquímica, Instituto de Química, Universidade de São Paulo (USP), São Paulo, SP, Brazil; Citrus Research and Education Center (CREC), Department of Microbiology and Cell Science, University of Florida, Lake Alfred, FL 33850, USA; Laboratório Nacional de Biociências, Centro Nacional de Pesquisa em Energia e Materiais, R. Giuseppe Máximo Scolfaro 10000, Campinas, SP, 13083-970, Brazil

**Keywords:** Interbacterial competition, Type IV secretions systems, CsrA, Single-cell fluorescent microscopy

## Abstract

Several *Xanthomonas* species have a type IV secretion system (T4SS) that injects a cocktail of antibacterial proteins into neighbouring Gram-negative bacteria, often leading to rapid lysis upon cell contact. This capability represents an obvious fitness benefit since it can eliminate competition while the liberated contents of the lysed bacteria could provide an increase in the local availability of nutrients. However, the production of this Mega Dalton-sized T4SS, with over a hundred subunits, also imposes a significant metabolic cost. Here we show that the chromosomal *virB* operon, which encodes the entirety of structural genes of the T4SS in *X. citri*, is regulated by the global regulator CsrA. Relieving CsrA repression from the *virB* operon produced a greater number of T4SSs in the cell envelope and an increased efficiency in contact dependent lysis of target cells. However, this was also accompanied by a physiological cost leading to reduced fitness when in co-culture with wild-type *X. citri*. We show that T4SS production is constitutive despite being downregulated by CsrA. Cells subjected to a wide range of rich and poor growth conditions maintain a constant density of T4SSs in the cell envelope and concomitant interbacterial competitiveness. These results show that CsrA provides a constant though partial repression on the *virB* operon, independent of the tested growth conditions, in this way controlling T4SS-related costs while at the same time maintaining *X. citri*’s aggressive posture when confronted by competitors.

**Author Summary:** *Xanthomonas citri* is a member of a family of phytopathogenic bacteria that can cause substantial losses in crops. At different stages of the infection cycle, these cells will encounter other bacterial species with whom they will have to compete for space and nutrients. One mechanism which improves a cell’s chance to survive these encounters is a type IV secretion system that transfers a cocktail of antimicrobial effector proteins into other Gram-negative bacteria in a contact-dependent manner. Here, we show that this system is constitutively produced at a low basal level, even during low nutrient conditions, despite representing a significant metabolic burden to the cell. The conserved global regulator, CsrA, provides a constant, nutrient-independent, repression on the production T4SS components, thereby holding production costs to a minimum while at the same time ensuring *X. citri*’s competitiveness during encounters with bacterial rivals.

## Introduction

Type IV secretion systems (T4SS) are large systems spanning both the inner and outer membrane of many Gram-negative bacterial species (Low et al. 2014). The best described functions of T4SSs are the delivery of the pTi plasmid of *Agrobacterium tumefaciens* (Vergunst et al. 2000; Gordon & Christie 2014), their crucial roles in bacterial conjugation (Ilangovan et al. 2015) and involvement in delivering virulence factors to mammalian cells by several pathogenic species (Dehio & Tsolis 2017). T4SSs are made up of over 100 subunits of 12 different proteins (VirB1 to VirB11 plus VirD4) with a total size of over 3 MDa each (Low et al. 2014). Maintaining and expressing large operons and producing the amino-acids required to assemble the proteins they encode present a significant investment in terms of energy and raw materials for a cell (Akashi & Gojobori 2002; Wagner 2005; Lynch & Marinov 2015). Given the high cost secretion systems would have on cell physiology, it is not surprising that their production is often restricted to specific conditions, where they will be most needed. For example, expression of the *Agrobacterium tumefaciens* T4SS is dependent on pH, monosaccharides, phosphate and specific phenolic compounds released by wounded plant tissue (Das & Pazour 1989; Lohrke et al. 2001; Winans 1990; Gao & Lynn 2005).

Similarly, plasmid-borne *tra* genes, encoding a T4SS and other components of the conjugation machinery, are only produced during specific conditions and often only in a small portion of the population (Koraimann & Wagner 2014). Other examples include the *Brucella suis* T4SS produced inside acidic phagocytic vacuoles of macrophages (Boschiroli et al. 2002) and the *Ehrlichia ruminantium* T4SS whose genes are induced during iron starvation (Moumène et al. 2017). This strict environmentally-dependent regulation is also common in other secretion systems; for example, the *Vibrio cholera* type VI secretion system (T6SS) is induced during high cell densities on chitinous surfaces (Borgeaud et al. 2015), the *Xanthomonas citri* T6SS is specifically induced in the presence of amoeba (Bayer-Santos et al. 2018) and the *Shigella flexneri* type III secretion system is tightly regulated by oxygen levels (Marteyn et al. 2010).

*X. citri* is a phytopathogen that causes citrus canker, a disease which can lead to significant losses in citrus fruit production (Ryan et al. 2011; Mansfield et al. 2012). Previously, our group has characterized the interbacterial killing activity of a T4SS in *X. citri* and showed that this strain actively transfers a cocktail of antibacterial effector proteins into neighbouring Gram-negative cells in a contact-dependent manner (Souza et al. 2015; Sgro et al. 2019). More recently, we also described the antibacterial killing of a similar T4SS with its unique antibacterial effectors in the opportunistic pathogen *Stenotrophomonas maltophilia* (Bayer-Santos et al. 2019). Despite the importance of T4SSs in interbacterial competition, little is known concerning the regulation of these systems in Xanthomonadaceae.

Microarray data of an *X. citri* strain harbouring a mutation in the global regulator CsrA (also called RsmA) indicated its involvement in the regulation of over a hundred genes, including the *virB* operon that encodes the T4SS proteins VirB1-11 (Andrade et al. 2014). CsrA is a pleiotropic regulator linked to the genetic changes during stationary phase growth, biofilm formation, gluconeogenesis and virulence (Romeo & Babitzke 2018). CsrA acts by binding specific mRNA loops in 5’ untranslated regions containing the canonical 5’-GGA-3’ motif (Liu & Romeo 1997; Holmqvist et al. 2016). In some cases, these interactions stabilize the mRNA leading to increased expression, as for example has been observed for the *hrpG* mRNA in *X. citri* (Andrade et al. 2014). More often, these CsrA-binding loops encompass the ribosome binding site, in which case binding of CsrA inhibits translation (Baker et al. 2002). Although several other means of CsrA regulation exist (Romeo & Babitzke 2018), the majority of interactions lead to a repression of protein production (Potts et al. 2017). CsrA is regulated by two important small RNAs, CsrB and CsrC (in *E. coli*) or by RsmY and RsmZ (in *P. aeruginosa*), which contain several high affinity CsrA binding loops that effectively titrate CsrA (Weilbacher et al. 2003; Janssen et al. 2018). Production of these small RNAs in *E. coli* is controlled by several regulatory pathways, including the BarA/UvrY two-component system that responds to molecules such as formate and acetate, the catabolite repression pathway mediated by cAMP-CRP and the stringent response governed by RelA and SpoT-mediated production of (p)ppGpp (Romeo & Babitzke 2018). Furthermore, direct regulation of CsrA copy numbers in *E. coli* is achieved by five different promoters using at least two different sigma factors (Yakhnin et al. 2011).

*X. citri* CsrA is very similar to CsrA from *E. coli* and *P. aeruginosa* (>77% identical), albeit the *X. citri* protein has a 9 residue C-terminal extension. Detailed knowledge of CsrA, its targets and its regulation in Xanthomonas species is limited. For example, the identity or presence of regulatory small RNAs is not known. Nonetheless, some studies have shown phenotypic alterations in a CsrA deletion strain reminiscent with known CsrA phenotypes in *E. coli* and *P. aeruginosa*, such as reduced virulence, increased biofilm formation and increased glycogen production (Chao et al. 2008; Lu et al. 2012) and direct RNA binding studies have also confirmed the affinity of CsrA in *X. citri* for the canonical 5’-GGA-3’ motifs (Andrade et al. 2014).

In this work, we show that the CsrA protein of *X. citri* represses the *virB* operon and that removal of CsrA repression has a measurable fitness cost. This repression is incomplete however, and so production of the *virB* products continues at a controlled basal level, maintaining a constant density of T4SSs in the cell envelope during different growth conditions. We propose that CsrA, in concert with other unidentified regulatory factors working on the *virB* operon, leads to a sustained and energetically affordable aggressive posture that contributes to *X. citri* competiveness and survival.

## Results

### CsrA regulates the virB operon by binding to the 5’ UTR of virB7

Based on transcription start site analysis data for *Xanthomonas campestris* (Alkhateeb et al. 2016) and further observations made in the Sequence Read Archive for *X. campestris and X. citri* (https://www.ncbi.nlm.nih.gov/sra), the *Xanthomonas vir* locus contains two main transcription start sites (TSSs) (Figure 1a). The presence of these TSSs in *X. citri* was confirmed by 5’RACE analysis (Supplemental Figure S1). The first TSS is located 303 nucleotides upstream of *virD4* and a second TSS is located 249 nucleotides upstream of the *virB7* start codon (Figure 1a and Supplemental Figure S1). Both *virD4* and *virB7* contain large upstream regions, with the upstream region of *virD4* most probably containing an open reading frame expressing a conserved protein of unknown function. Although an open reading frame can be detected in the region downstream of *virD4* and upstream of *virB7* (nucleotide sequence shown in Figure 1a), it lacks a canonical ribosome binding site and its translated product is of very low sequence complexity and is not conserved in the protein databases. Since a list of CsrA regulated genes in *X. citri* from a microarray dataset included several of the *virB* genes (Andrade et al. 2014) we further scrutinised the *virB* 5’UTR (upstream of the virB7 start codon, from here on referred to as *5’UTR*_*B7*_). This analysis revealed several 5’-GGA-3’ CsrA-binding motifs (Figure 1a) that could be CsrA binding sites when present in a stable loop structure (Holmqvist et al. 2016). To test this hypothesis, a transcriptional *msfgfp* (encoding for the monomeric super folder green fluorescent protein) fusion was constructed downstream of the genomic copy of *virB11* (*X. citri virB11-msfgfp*). Single-cell fluorescence analysis showed that upon deleting *csrA* in *X. citri virB11-msfgfp,* msfGFP production from the *virB* operon is upregulated 3.3-fold (Figure 1b). In order to confirm the direct regulation of CsrA on the *5’UTR*_*B7*_, the *5’UTR*_*B7*_ was cloned in between the P_*tac*_ promotor (constitutive in the *lac* negative *X. citri* strain) and *msfgfp,* after which a single copy of this construct was integrated into the α-amylase gene (*amy; xac0798*) in both the *X. citri* wild-type and Δ*csrA* strain. The resulting fluorescence levels in the presence or absence of CsrA shows that CsrA is capable of repressing msfGFP production from the P_*tac*_ promoter when the *5’UTR*_*B7*_ is present (Figure 1c). This indicates that the effect of CsrA is independent of the *virB* promoter. Expanding these observations, we replaced the entire structural operon (Figure 1a) by *msfgfp,* with the *msfgfp* start codon in the exact position of the *virB7* start codon (*X. citri ∆virB::P*_*B7*_*-msfgfp*). Deleting *csrA* in this strain indeed showed a similar response of increased msfGFP production from the *virB* operon (Figure 1d). Importantly, the removal of the *5’UTR*_*B7*_ in this strain (*X. citri ∆virB::P*_*B7*_*-∆5’UTR*_*B7*_*-msfgfp*) caused msfGFP levels to increase in the wild-type strain (Figure 1d). This increase in msfGFP production in the absence of *5’UTR*_*B7*_ was only slightly lower than those observed in the *X. citri ∆virB::P*_*B7*_*-msfgfp ∆csrA* and *X. citri ∆virB::P*_*B7*_*-∆5’UTR*_*B7*_*-msfgfp ∆csrA* strains, further confirming CsrA regulation mediated by the 5’UTR_B7_. We note that the *∆csrA* background has an elaborate effect on *X. citri* physiology (for instance, cultures display an intense flocculation and decreased cell-sizes) and that this could account for the subtle differences in msfGFP production observed between strains lacking *csrA* with or without the *5’UTR*_*B7*_ in Figure 1d. Next, the construction of a genomic deletion of the *5’UTR*_*B7*_, while keeping all other *virB* genes intact, resulted in a 3.9-fold increase in expression levels in the *X. citri ∆5’UTR*_*B7*_ *virB11-msfgfp* reporter strain (Figure 2a). Finally, an electrophoretic mobility shift assay (EMSA) confirmed the direct *in vitro* binding of CsrA to the *5’-UTR*_*B7*_ (Figure 1e). This result, combined with all the gene expression data obtained from several different constructs (Figures 1b to 1d and 2a) indicate that CsrA regulates *virB* operon expression by binding to the *5’UTR*_*B7*_, most probably by preventing translation and/or destabilizing the *virB* transcripts.

**Figure 1:**
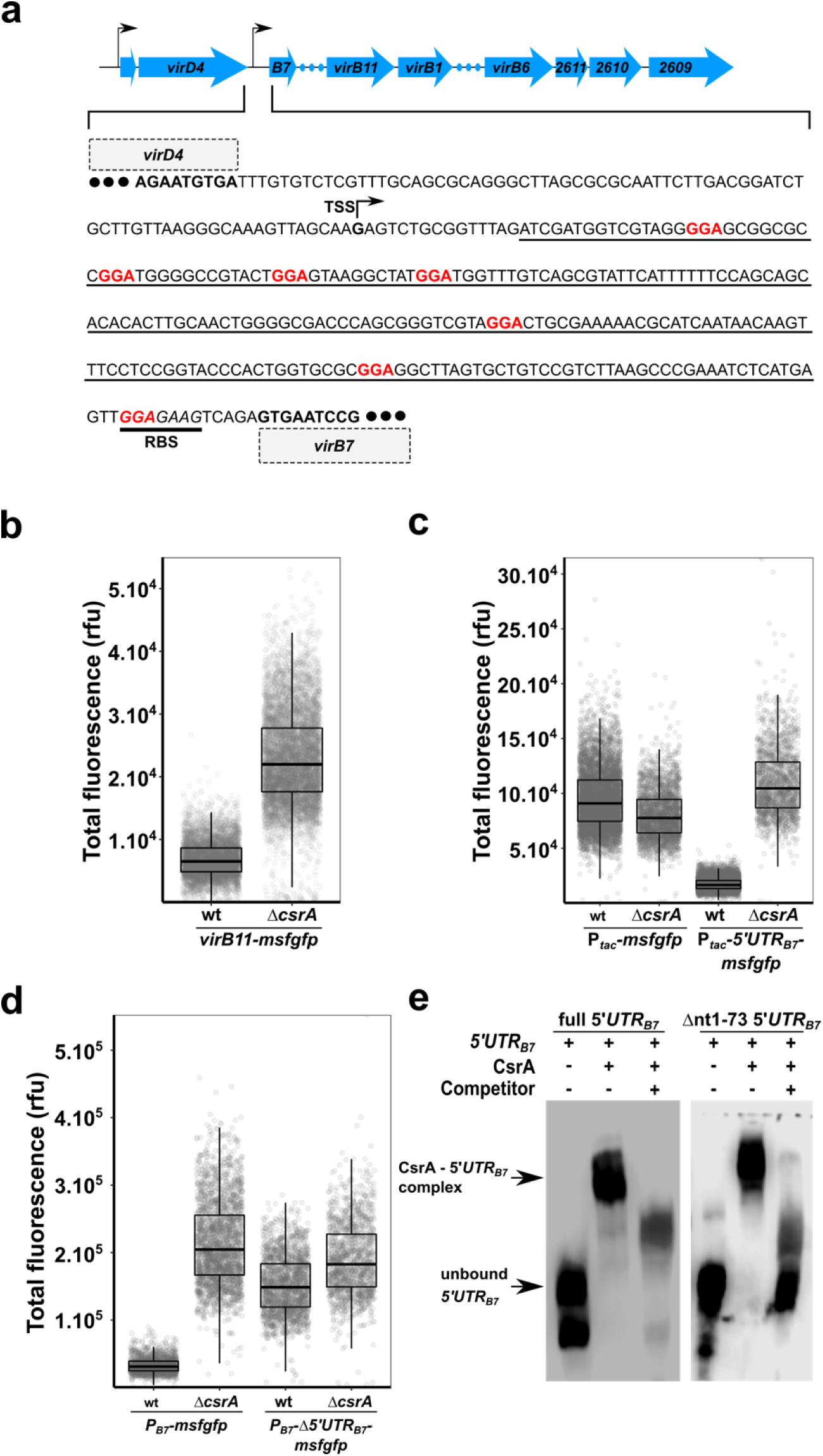
CsrA regulates the *virB* operon. **a)** An on-scale schematic of the *virD* and *virB* operons with black arrows indicating positions of the transcription start sites, blue arrows indicating the open reading frames and blue dots representing genes not shown. The intergenic sequence between *virD4* and *virB7* genes is shown and the *virB7* transcription start site (TSS) is indicated. The 5’-GGA-3’ motifs are indicated in red and the predicted ribosome binding site (RBS) is underlined with a thick black line. The sequence that is deleted in the *∆5’UTR*_*B7*_ strains is underlined with a thin black line. **b)** Single-cell msfGFP fluorescence levels of the *X. citri virB11-msfgfp* transcriptional reporter in *X. citri* wild type and *∆csrA* strains. Production of msfGFP increased on average 3.3-fold in the *∆csrA* strain (N_wt_ = 4887 cells and N_∆csrA_ = 6263 cells, from a single representative culture each). **c)** Single-cell msfGFP fluorescence levels of the *amy::P*_*tac*_*-msfgfp* and *amy::P*_*tac*_*-5’UTR*_*B7*_*-msfgfp* reporters in *X. citri* wild type and *∆csrA* strains. A CsrA-dependent reduction in msfGFP production is only observed in the strain containing the *5’UTR*_*B7*_ (average 5.8-fold reduction, N_wt_ = 6862 cells and N_∆csrA_ = 1671 cells, from 2 separate cultures each). The control lacking the *5’UTR*_*B7*_ in between the P_*tac*_ promotor and *msfgfp* (*X.citri amy::P*_*tac*_*-msfgfp*) showed no CsrA-dependent repression of msfGFP production (average 16% increase, N_wt_ = 5534 cells and N_∆csrA_ = 2495 cells, from 2 separate cultures each). **d)** Single-cell msfGFP fluorescence levels of the *msfgfp* reporter that substituted the entire structural *virB* operon (with the *msfgfp* start codon placed at the exact position of the *virB7* start codon) in *X. citri* wild type and *∆csrA* strains with an intact *virB* promoter (*P*_*B7*_) or a *virB* promoter in which the *5’UTR*_*B7*_ was deleted (*P*_*B7*_*-∆5’UTR*_*B7*_). *X. citri ∆virB::P*_*B7*_*-∆5’UTR*_*B7*_*-msfgfp* shows higher msfGFP production than the *X. citri ∆virB::P*_*B7*_*-msfgfp* strain containing the wild type 5’UTR_B7_ (average 6.5-fold increase, N_wt_ = 1495 cells and N_∆csrA_ = 2156 cells, from 2 separate cultures each). Absence of the *5’UTR*_*B7*_ almost completely abolishes CsrA dependent down-regulation of expression levels (average 1.2-fold increase, N_wt_ = 1748 cells and N_∆csrA_ = 1304 cells, from 2 separate cultures each). Note that removal of the *5’UTR*_*B7*_ in the wild-type backgrounds led to a 4.6-fold increase. **e)** An electrophoresis mobility shift assay (EMSA) shows direct *in vitro* binding of CsrA with the complete and a shortened fragment (lacking the first 73 nucleotides) of the *5’UTR*_*B7*_. Addition of unlabelled *5’UTR*_*B7*_ RNA competes with labelled RNA binding. Tukey box-and-whisker plots in parts b, c and d: black central line (median), box (first and third quartiles) and whiskers (data within 1.5 interquartile range).

**Figure 2:**
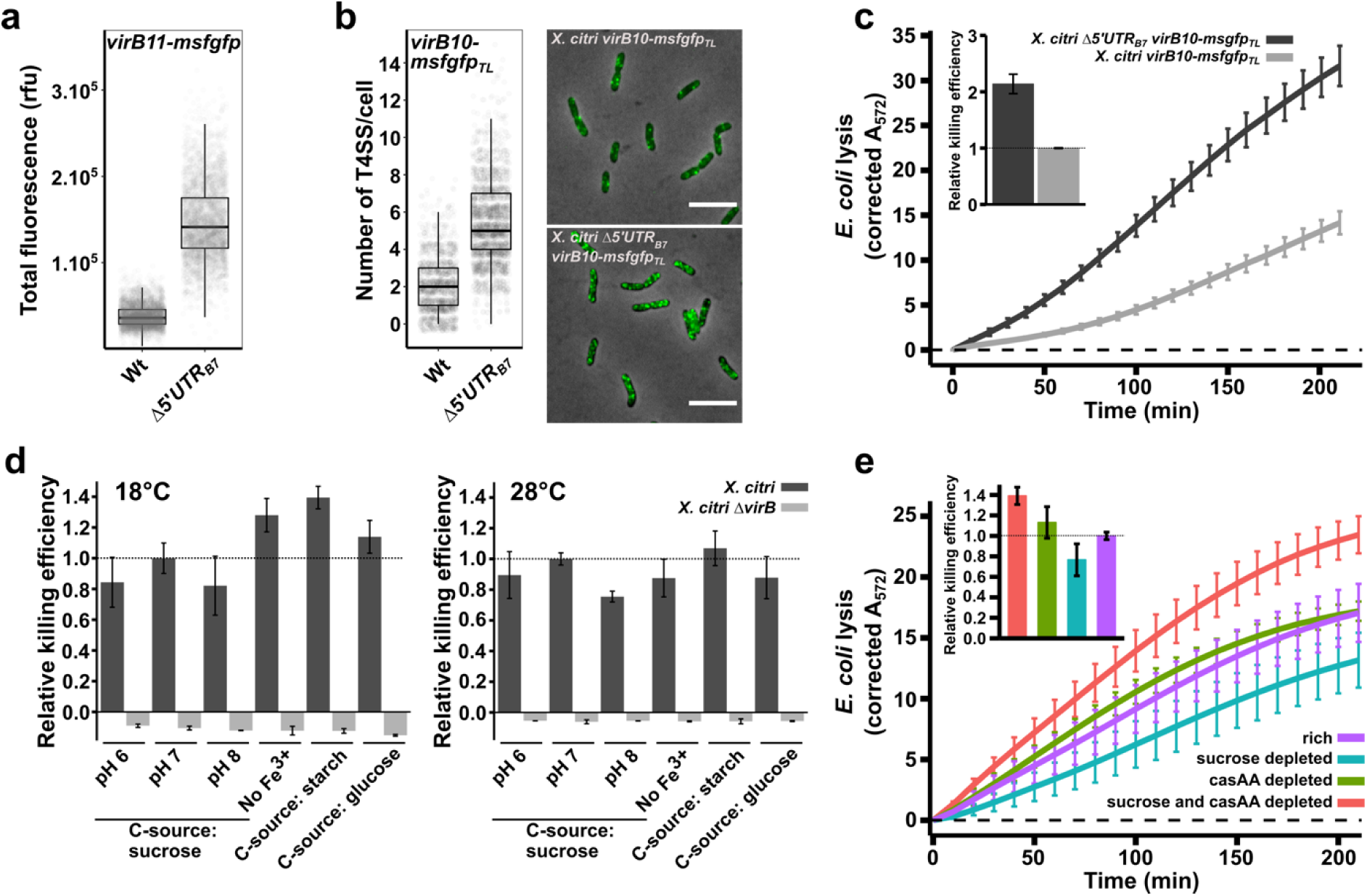
VirB production and interbacterial killing is constitutive under incomplete repression by CsrA. **a)** Single-cell msfGFP fluorescence levels from the *X. citri virB11-msfgfp* reporter strain with and without the genomic deletion of the *5’UTR*_*B7*_ (underlined region in Figure 1a; *X. citri ∆5’UTR*_*B7*_ *virB11-msfgfp*). Removing the *5’UTR*_*B7*_ leads to a 3.9-fold increase in msfGFP production from the *virB* operon (N_wt_ = 3397 cells and N_∆5’UTR_ = 3099 cells, from 4 separate cultures). **b)** Quantification of single fluorescent foci of T4SSs containing the VirB10-msfGFP chimera. The genomic deletion of the *5’UTR*_*B7*_ in the *X. citri virB10-msfgfp*_*TL*_ strain increased the number of T4SSs per cell 2.6-fold (N_wt_ = 1361 cells and N_∆5’UTR_ = 2062 cells, from 3 separate cultures). Note that the increased amount of T4SS in the *∆5’UTR*_*B7*_ strain in some cases causes the VirB10-msfGFP produced fluorescent foci to overlap, limiting the separation of the single foci, leading to an underestimation of the number of T4SS per cell. **c)** Quantitative CPRG cleavage-based killing assay in the presence or absence of CsrA repression on the *virB* operon using the same *X. citri virB10-msfgfp*_*TL*_ and *X. citri ∆5’UTR*_*B7*_ *virB10-msfgfp*_TL_ cultures analysed in part **b**. Genomic deletion of the *5’UTR*_*B7*_ leads to a 2.14-fold increase in lysis of *E. coli* cells. The inset shows the slope value of the linear part of the depicted curves relative to the wild-type strains (N = 4 separate cultures with 2 technical repeats each). **d)** Quantitative CPRG cleavage-based killing assays of wild type *X. citri* cells grown in different conditions (dark grey bars). Killing efficiencies were evaluated in defined media containing different carbohydrate sources (0.2 % sucrose, 50 µg/ml starch or 0.2% glucose), pH 6, 7 and 8, lack of Fe^3+^ and at different temperatures (18°C and 28°C). Values represent the slope of the linear part of the OD_572_ curves as described in panel c and normalized relative to the condition at pH 7 (which is the condition used throughout the manuscript). As a control, killing efficiency of a T4SS-deficient mutant (*X. citri ∆virB::P*_*B7*_*-msfgfp*; light grey bar) was assessed after growth under the same conditions (N = 4 separate cultures with two technical repeats). **e)** Quantitative CPRG cleavage-based killing assays from *X. citri* cells grown at different nutrient levels. Killing efficiencies of wild type *X. citri* strains are maintained during growth in defined media containing either 0.2% or 0.01% sucrose and/or casamino acids (casAA). Inset shows the slope value of the linear part of the depicted curves relative to the killing curve from the *X. citri* cultures grown in rich media. Cultures reduced in both casamino acid and sucrose concentrations show a 39% increase whereas sucrose depleted cultures have a 23% reduction in efficiencies compared to the reference (N = 5 separate cultures with 4 technical repeats). Error bars in panels c, d and e indicate the standard deviation. Horizontal dashed line in c and e represents the zero-line obtained after subtracting the background signal of non-lysed *E. coli* cultures grown in parallel during each experiment.

### Removal of CsrA repression on the virB operon increases T4SS numbers and bacterial killing

Having established the direct role of CsrA in the repression of the *virB* operon, we asked whether the interbacterial killing efficiency of *X. citri* would be enhanced in a strain lacking the *5’UTR*_*B7*_. We used a *X. citri virB10-msfgfp*_*TL*_ translational fusion strain (Sgro et al. 2018) in order to assess the number of T4SSs that are present per cell (see Materials and Methods and Figure 2b). In this strain, the periplasmic VirB10 component has been replaced by a VirB10-msfGFP chimera. Since each T4SS contains 14 copies of VirB10, assembled T4SSs can be observed as fluorescent periplasmic foci and counted (Sgro et al. 2018). Deleting the *5’UTR*_*B7*_ in the *X. citri virB10-msfgfp*_*TL*_ genetic background resulted in a 2.6-fold increase in the number of fluorescent T4SS foci that were counted per cell (Figure 2b). We note that the higher density of T4SSs in this strain leads to a more crowded periplasm, making it more difficult to clearly separate individual foci, leading to an underestimation of total number of T4SSs. Therefore, the calculated 2.6-fold increase should be considered a lower limit. As could be expected, a higher number of T4SSs also increased the efficiency with which *X. citri* lyses *E. coli* cells in a quantitative LacZ mediated CPRG-cleavage assay (Figure 2c). Using the slopes of the curves in the CPRG-cleavage assays as a measure of killing efficiency (see Materials and Methods and (Sgro et al. 2018)), the *Δ5’UTR*_*B7*_ strain kills 2.14-fold more efficiently than the *X. citri* wild-type strain under these conditions (Figure 2c). However, these results also show that under these conditions, removal of CsrA repression was not necessary to observe T4SS dependent *E. coli* lysis in the CPRG assays (Figure 2c). This is in agreement with previously published spot assays and CFU-based competition assays, all performed with wild type *X. citri* strains (Souza et al. 2015). To test whether we could identify conditions in which T4SS-mediated killing would be inhibited or enhanced, we tested *E. coli* lysis efficiencies at 18°C or 28°C, at pH 6, 7 and 8, in absence of Fe^3+^ and using different carbohydrate sources (glucose, sucrose or starch; Figure 2d). All the tested conditions led to clearly detectable T4SS dependent lysis of *E. coli* cells with only small variations (+/- 35%) in killing efficiencies (Figure 2d). Taken together, these observations indicate that the observed levels of msfGFP signal from the *X. citri virB11-msfgfp* reporter strain (Figure 1b, 1d and 2a) and the discrete numbers of T4SSs in the cell envelope (Figure 2b) present in the wild-type background, represent the baseline expression and production levels of T4SS components and that these levels, all repressed from what they would otherwise be in the absence of CsrA, are sufficient to maintain T4SS production and efficient lysis of neighbouring target cells.

### T4SS dependent interbacterial killing is maintained during growth under scarce nutrient conditions

Considering the expected high energetic cost of producing T4SSs and the previously described responsiveness of *csrA* regulons to nutrient input (Romeo & Babitzke 2018), we decided to test whether lowering the nutrient contents of the growth media would change T4SS-dependent killing efficiencies. To accomplish this, *X. citri* cultures were passed to media containing normal or reduced levels of sucrose and/or casamino acids (the only nutrient sources present in the defined media) and grown overnight plus an additional 9 hours in fresh media for each culture to fully adapt to the conditions (see Materials and Methods for details). These cultures were then subjected to a quantitative CPRG-cleavage assay under standardized conditions. Figure 2e shows that *X. citri* grown in nutrient-scarce culture media continues to sustain T4SS-dependent lysis of *E. coli* cells. *X. citri* cultures with limited access to sucrose have slightly reduced killing efficiencies (23% decrease) while cells with limited access to both casamino acids and sucrose presented slightly elevated (39% increase) killing efficiencies (Figure 2e). Supplemental Movie S1 presents a time-lapse video of a mixed culture of *X. citri* and *E. coli* cells growing in casamino acid-depleted media for over 30 hours in which many events of *E. coli* cell lysis are observed upon contact with *X. citri* cells. For comparison, Supplemental Movie S2 presents *X. citri* killing *E. coli* cells growing under standard nutrient concentrations and also shows clear cell lysis upon contact with *X. citri* cells. No *E. coli* lysis is observed in these experiments when T4SS-deficient *X. citri* strains are employed ((Souza et al. 2015) and data not shown).

### Constant but incomplete CsrA-mediated repression of the virB operon during different growth conditions

Figures 2a and 2b, showed that removing the *5’UTR*_*B7*_ leads to both increased msfGFP production and T4SS assembly. In order to test how CsrA regulation impacts protein production from the *virB* operon, we decided to look more closely at VirB production in the presence and absence of the *5’UTR*_*B7*_ under different growth conditions. To do this, we began by comparing cytoplasmic msfGFP production from *X. citri virB11-msfgfp* and *X. citri ∆5’UTR*_*B7*_ *virB11-msfgfp* strains. Both strains were grown simultaneously in media with sucrose, glucose, glycerol or starch as carbohydrate sources or in media depleted for casamino acids and/or sucrose. Cultures were grown overnight in the different media and after diluting, grown for an additional 6 hours and msfGFP content was registered by fluorescence microscopy.

Mean msfGFP production from the *X. citri virB11-msfgfp* strain remains fairly similar during different inputs of carbohydrates and even in the absence of a carbohydrate source (Figure 3a; white boxplots). Reducing casamino acid concentrations, however, increases VirB production 60% on average (both in combination with 0.2% or 0.01% sucrose). This increase in expression levels seems to be independent of CsrA regulation since similar increases in msfGFP production were detected when using the *X. citri ∆5’UTR*_*B7*_ *virB11-msfgfp* strain (Figure 3a, dark grey boxplots). In fact, when comparing the relative increases observed with the removal of the *5’UTR*_*B7*_, it appears that repression on the *5’UTR*_*B7*_ leads to a reduction of VirB protein production by, on average 3.8-fold (± 0.4) over all conditions tested for both strains (Figure 3a, fold-changes are indicated above each pair of boxplots). As such, it appears that CsrA represses the *virB* operon to an extent that is largely independent of the carbohydrate source or casamino acid availability.

**Figure 3:**
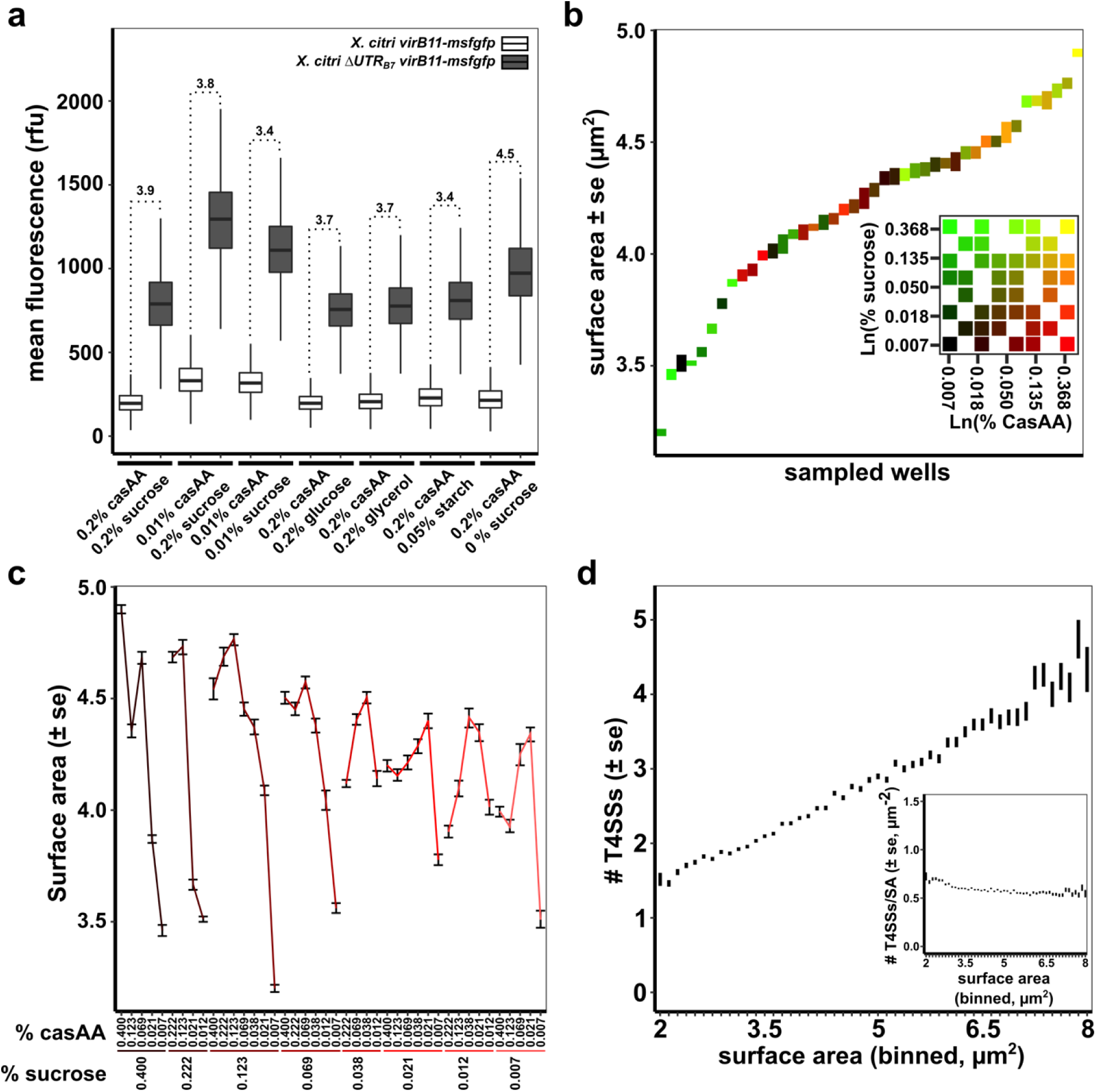
T4SS production and density in the cell envelope under different nutrient conditions. **a)** msfGFP fluorescence levels from *X. citri virB11-msfgfp* and *X. citri ∆5’UTR*_*B7*_ *virB11-msfgfp* grown in AB defined media containing either sucrose, glucose, glycerol, starch or no carbohydrate source (0% sucrose) in combination with 0.2% casamino acids (casAA) or in AB defined media containing 0.01% casamino acids with 0.2% or 0.01% sucrose. Mean cytoplasmic msfGFP production from the *X. citri virB11-msfgfp* transcriptional fusion strain in the presence (white boxplots) or absence (grey boxplots) of the *5’UTR*_*B7*_ are represented as Tukey box-and-whisker plots for each media composition with black central line (median), box (first and third quartiles) and whiskers (data within 1.5 interquartile range). On average 6,018 cells were sampled per condition for each strain, with a minimum of 2,162 and a maximum of 16,351 cells from two independent cultures. Note that in this graph, the mean fluorescence (sum of each cell’s pixel intensities divided by the number of pixels) is used to take into account differences in cell-sizes between conditions. **b)** Surface area of *X. citri* cells sampled from separate cultures, each culture differing slightly in sucrose and casamino acid concentrations. Cultures are ordered according to the average cell surface area observed for each growth condition. The inset shows the 8×8 matrix range of sucrose and casamino acid concentrations used as described in the text. In both the graph and the inset, each growth condition is colored differently. **c)** Surface area of *X. citri* cells as a function of sucrose and casamino acid concentrations. The data is the same as in panel b but organised with respect to sucrose and casamino acid concentrations and illustrates the link between *X. citri* cell sizes and the culture media. **d)** Average number of T4SSs per cell versus cell surface area from all the cells grown in the different conditions presented in panels b and c. Detection of VirB10-msfGFP foci reveals that the increase of average number of T4SSs correlates with increasing surface area (Pearson correlation r = 0.34, as calculated for all data points). *Inset:* Density of T4SSs in the cell envelope (number of T4SS per surface area, T4SS/SA) versus the surface area. A subtle increase with smaller (and thus more nutrient deprived) cells is observed. Vertical bars in figure b, c and d represent the standard error around the mean (± se). In total 69,412 cells were registered for the data in panels b, c and d.

### Homeostasis of T4SS density in the cell envelope over a wide range of nutrient availability

Spurred by the increased expression of the *virB* operon during decreased inputs of casamino acids, we set out to assess the number of assembled T4SS over a wide range of casamino acid and sucrose concentrations. For this, an 8 by 8 matrix of wells containing defined media with varying nutrient levels was created in a 96-well plate. With sucrose (rows) and casamino acids (columns) ranging from 0.4% to 0.007% in a 1.8x dilution series. After 24 hours of growth and an additional 5 hours of growth after a 4-fold dilution in the same but fresh media, *X. citri virB10-msfgfp*_*TL*_ cells were sampled and immediately imaged by fluorescence microscopy, registering both cell dimensions and the number of T4SS foci present per cell. In total, 42 different conditions (Figure 3b inset) were sampled over two separate experiments). Figures 3b and 3c show that the different nutritional inputs in each well result in a range of cell sizes, represented by their average surface areas and reveal a general trend of reduced cell size with reduced casamino acid or sucrose availability. Plotting the number of T4SSs versus the surface area shows a linear increase in T4SS numbers with increasing surface area (Figure 3d; Pearson correlation *r* = 0.34, as calculated for all data points). However, this positive correlation in turn leads to an almost constant average T4SS density (T4SS/surface area) in the cell envelope with a subtle increase observed for the smallest cells (Figure 3d inset), in line with the observation of subtly increased msfGFP production in the *X. citri virB11-msfGFP* transcriptional reporter at low casamino acid concentrations (Figure 3a). These results also suggest that during the cell-cycle, when the surface area gradually increases until cell division, T4SSs are added continuously. Therefore, it seems that a *X. citri* population maintains the density of its T4SSs within a specific range under a variety of nutritional conditions. The number of T4SSs relative to surface area (T4SS density) could be an important factor in determining the probability that a *X. citri* cell is able to successfully transfer effectors into a neighbouring target cell during interbacterial competition.

### T4SS overproduction in a ∆5’UTRB7 background has an impact on X. citri physiology and leads to reduced growth speeds

Since *∆5’UTR*_*B7*_ cells present a roughly 4-fold greater expression from the *virB* operon (Figure 3a) and kills with approximately twice the efficiency as wild-type cells (Fig. 2c), we asked whether this putative advantageous feature could be counter-balanced by the inherent metabolic cost of T4SS production. We therefore set up a co-culture experiment to test whether overproduction of T4SSs in the *∆5’UTR*_*B7*_ background leads to a detectable growth defect in *X. citri*. For this we took advantage of the difference in msfGFP production levels between *X. citri virB11-msfgfp* and *X. citri ∆5’UTR*_*B7*_ *virB11-msfgfp* (Figures 2a and 3a) to sort wild type and overproduction cells by fluorescence microscopy. This mitigated the need to introduce different antibiotic resistance markers in the genome that could on their own lead to subtle physiological differences. Additionally, the strains used here are genetically very closely related since they were obtained from the same recombination events leading to either the wild-type or mutant *5’UTR*_*B7*_ allele (see Materials and Methods). Separate cultures of different single colonies of each of the strains were synchronised in rich AB media before being mixed and diluted in a 1:1 ratio into AB media supplemented with 0.2% sucrose and 0.01% casamino acids. Cultures were diluted regularly to avoid saturation and fluorescence microscopy images of thousands of cells were obtained at different time-points over a period of approximately one week. Figure 4a shows that the *∆5’UTR*_*B7*_ strain gets outpaced by the wild type *5’UTR*_*B7*_ strain. In two distinct experiments using 3 or 4 separate cultures each, we observed a 19% and 13% average decrease in the *X. citri ∆5’UTR*_*B7*_ *virB11-msfgfp* cell population relative to the wild-type *X. citri virB11-msfgfp* cell population (Figure 4a). Additionally, analysis of the cell sizes of the two strains at several time points over a one-week period revealed that the *∆5’UTR*_*B7*_ background consistently has a slightly larger surface area over volume ratio (SA/V; Figure 4b), indicative of smaller cell sizes. The SA/V ratio has been suggested to be a measure of the physiological state of rod-shaped bacterial cells and is thought to be set by the availability of nutrients (Harris & Theriot 2016; Harris & Theriot 2018). The observed average size reduction (increase in SA/V) for *X. citri ∆5’UTR*_*B7*_ *virB11-msfgfp* cells could be attributed to the increased metabolic cost of nutrients consumed in T4SS production and/or to stress induced by the greater number of T4SSs in the cell envelope. This experiment reveals that T4SS production has a measurable cost and illustrates the importance of balancing expenses with gains from increased aggressiveness during inter-bacterial competition.

**Figure 4:**
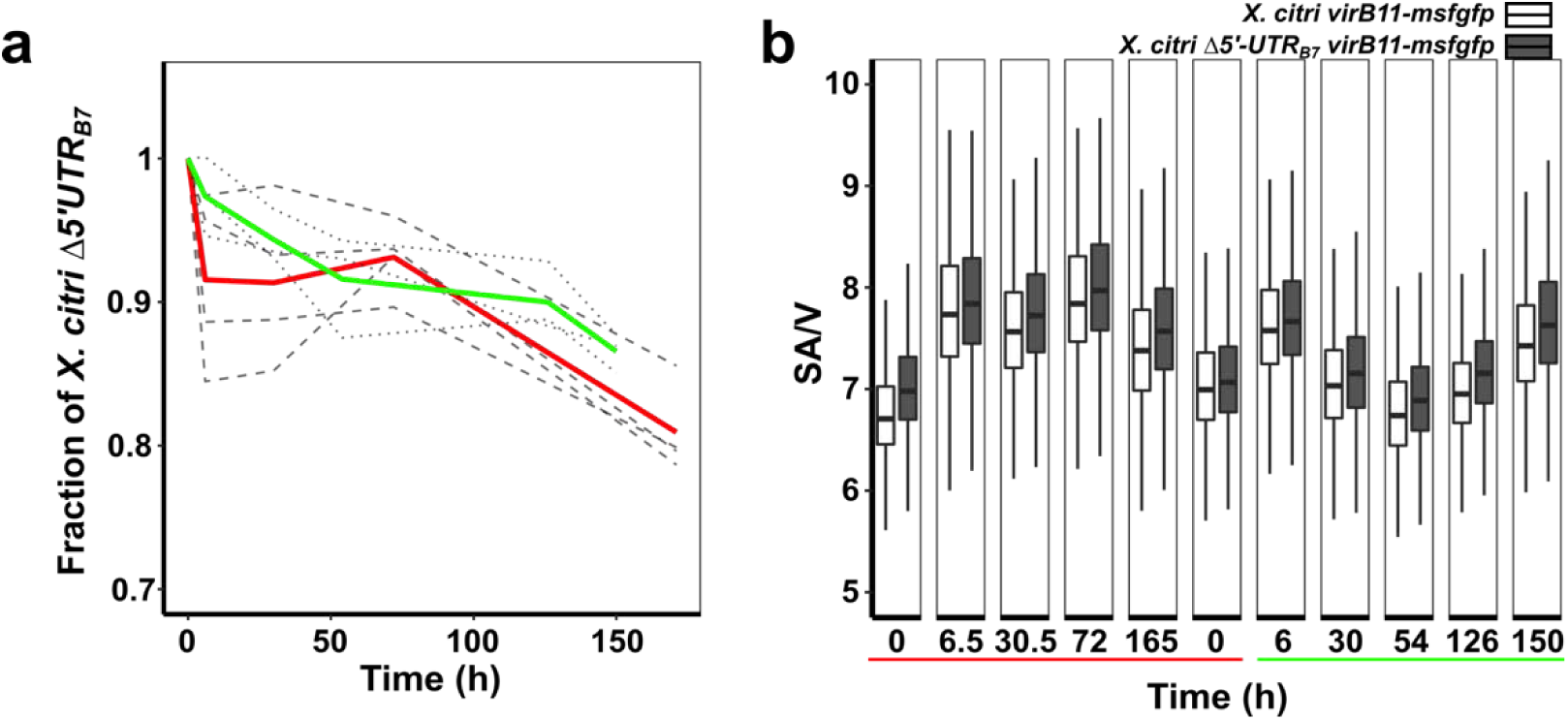
Overproducing T4SSs leads to a detectable physiological cost for *X. citri*. **a)** Co-culture experiment between *X. citri virB11-msfGFP* and X. *citri ∆5’UTR*_*B7*_ *virB11-msfGFP* in liquid medium. A decrease in the fraction of *X. citri ∆5’UTR*_*B7*_ *virB11-msfGFP* cells relative to the wild-type (*X. citri virB11-msfGFP*) cells is observed and becomes pronounced after approximately 1-week incubation. Results from two distinct experiments with 3 (dotted lines and green average) and 4 (dashed lines and red average) separate cultures are shown. In all seven cultures, we observed a significant reduction in the fraction of *X. citri ∆5’UTR*_*B7*_ *virB11-msfgfp* cells relative to the wild-type *X. citri virB11-msfgfp* cells at the end-points: 0.87 ± 0.01 (green) and 0.81 ± 0.03 (red). Cells were imaged by fluorescence microscopy and separated by the difference in msfGFP production levels as seen in Figures 2a and 3a. **b)** Comparison of surface area to volume ratios of *X. citri ∆5’UTR*_*B7*_ *virB11-msfGFP* and wild-type cells for cells collected at different times for all seven cultures shown in part a. A subtle but consistently smaller average cell size (larger SA/V) in the *∆5’UTR*_*B7*_ background is observed indicative of a subtle change in physiological state of the cells that can be attributed to the increased production of T4SSs. At each time point an average of 3871 cells were analysed for each culture (minimum: 1061 cells, maximum: 9582 cells).

## Discussion

We show here that the regulation of the *virB* operon in *X. citri*, coding for the structural proteins of the T4SS, is under control of the global regulator CsrA which binds to sites in the 5’ untranslated region of the polycistronic mRNA initiating at the *virB7* gene (*5’UTR*_*B7*_; Figure 1). The removal of the *5’UTR*_*B7*_ leads to an increase in production of *virB* encoded proteins, which in turn leads to an increase in number of T4SSs present in the cell envelope and a greater interbacterial killing efficiency (Figure 2a, 2b and 2c). Despite CsrA repression on the *virB* operon, we have not yet found any growth condition that would greatly decrease nor increase T4SS production and T4SS-dependent killing by wild-type *X. citri* cells (Figures 2d, 2e and 3a). Therefore, the removal of CsrA repression does not seem to be required to induce production of the T4SS. Rather, CsrA repression maintains a discrete number of T4SSs in the cell envelope over a range of different growth conditions tested (Figures 2 and 3).

Several factors can be imagined influencing the efficiency with which a *X. citri* cell can mount a successful contact-dependent attack. Firstly, a T4SS needs to be present at the contact interface between the attacking *X. citri* cell and the target rival cell. After a successful contact, T4SS effectors need to be translocated through the T4SS. This is dependent on both the availability of effectors and importantly, ATP to power the secretion. For example, the subtle decrease in killing efficiencies in the sucrose-depleted conditions (Figure 2e) might stem from reduced energy levels, since sucrose would be the main carbohydrate fed into the glycolysis and citric acid metabolic pathways. Furthermore, depleting nutrients also leads to much smaller cell sizes compared to the cells grown in rich media (Harris & Theriot 2016). This reduction in cell size in itself could influence killing-efficiencies by increasing the probability of small cells contacting larger *E. coli* cells when in co-culture on a solid surface, since smaller *X. citri* cells will have a more close-packed arrangement next to *E. coli* cells (Hudson 1949), increasing the probabilities of successful T4SS contact. Expanding this line of reasoning, a unit mass of cells formed during growth in reduced nutrient conditions would have a higher contact probability as the same mass of cells formed during growth in rich conditions, because of the former’s greater number of single cells. Thus, the subtle increase in T4SS density with lower nutrient inputs (Figure 3b) and the concomitant smaller cell sizes, could both be responsible for the slight increases in killing efficiency in during growth in media depleted in both casamino acids and sucrose (Figure 2e).

The complex regulatory features governing CsrA production and activity have been shown to be orchestrated in response to a wide variety of signals (Yakhnin et al. 2011; Romeo & Babitzke 2018). Complex regulation mechanisms, including negative autoregulation, have been proposed to act to keep CsrA levels and activity relatively stable with greatly reduced cell-to-cell variability, making CsrA an ideal regulator for homeostatic responses (Yakhnin et al. 2011; Romeo & Babitzke 2018). Of note, an ancestral CsrA homolog has been shown to act as a homeostatic control agent during flagella morphogenesis in the Gram positive *Bacillus subtilis* (Mukherjee et al. 2011). As such, CsrA could integrate diverse signals that relay information regarding the nutritional environment of the cell and subsequently stabilise its own activity to ensure a constant regulatory effect on its target transcripts. In light of this model of CsrA function, our experiments do indeed indicate that in *X. citri,* CsrA acts to reduce production from the *virB* operon with roughly the same repressing power over a wide range of growth conditions (Figure 3a). Recently, it was reported that CsrA (RsmA) represses all three Type VI secretion systems in *Pseudomonas aeruginosa*, abolishing translation under non-inducing conditions (Allsopp et al. 2017). Affinity of RNA loops for CsrA can differ in several orders of magnitude (Duss et al. 2014) and so some CsrA-mRNA associations will be very sensitive to fluctuations in mRNA levels and CsrA availability while others remain insensitive. As such, CsrA–mRNA affinities could be tuned so as to constitutively stabilise translation from one transcript at basal levels (such as for the *virB* operon) and at the same time ensure that translation from other transcripts is only derepressed by a specific trigger; for example by a specific condition that changes the mRNA structure. Several genetic circuits leading to different outcomes are possible, but for sake of general discussion, we note that it is unlikely that CsrA availability varies greatly under different conditions, since this would be hard to reconcile with the simultaneous control of the hundreds of transcripts through which CsrA exerts its pleiotropic effects (Andrade et al. 2014; Potts et al. 2017).

The removal CsrA-based repression of T4SS production results in a two-fold increase in interspecies bacteria killing efficiency (Figure 2c) which could be beneficial under certain circumstances. However, the production of a T4SS has its costs: the maintenance and transcription of an approximately 13 kb locus, multiple rounds of translation to produce the over 100 subunits that need to be transported to the inner-membrane or periplasm and assembled into a single system of over 3 MDa in size (Low et al. 2014). Since the maintenance of a single gene and the production of the amino acids to build up its protein product has a measurable cost (Akashi & Gojobori 2002; Wagner 2005; Lynch & Marinov 2015), it was not surprising that we were able detect a reduction in the fitness of the *X. citri* strain in which CsrA-based repression was removed (Figure 4).

Inter-bacterial competition is increasingly being shown to be crucial for the fitness, survival (Lories et al. 2017; García-Bayona & Comstock 2018), structuring of bacterial populations (Nadell et al. 2016) and possibly contributing to bacterial evolution by lysis and subsequent uptake of DNA (Veening & Blokesch 2017). The continuous expression and activity of the T4SS under different growth conditions, further illustrates the importance of interbacterial killing and the benefits that are accompanied with it. The control of *X. citri* T4SS production by CsrA, seems to guarantee constant densities of T4SSs that provide protection against rival bacteria, but not too many to represent a metabolic burden, thus maintaining a balance likely to be crucial for *X. citri* in the varied natural environments it encounters during its life cycle, both within and outside of its plant host.

## Materials and Methods

### Bacterial strains, media and culturing

All strains used are listed in Supplemental Table S1. For all experiments, strains were grown in defined AB media containing 15mM (NH_4_)_2_SO_4_, 17mM Na_2_HPO_4_, 22mM KH_2_PO_4_, 50mM NaCl, 0.1 mM CaCl_2_, 1 mM MgCl_2_ and 3 µM FeCl_3_, at pH 7.0, supplemented with 10 µg/mL thiamine and 25 µg/mL uracil and varying concentrations of carbohydrate sources and casamino acids as described in the text. For cloning purposes standard lysogeny broth (5g/l yeast, 5 g/l NaCl, 10 g/l tryptone and 15 g/l agar) and 2xYT (5 g/l yeast, 10 g/l NaCl, 16 g/l tryptone and 15 g/l agar) were used. For counterselection, sucrose plates were used (5 g/l yeast, 10 g/l tryptone, 60 g/l sucrose). Standard incubations were performed at 28°C in 24-well plates using 1.5 ml culture media or in 96-well plates with 200 µl culture media with shaking at 200 rpm. In general, after a first overnight growth period in 2xYT medium, cells were transferred at a 100-fold dilution into AB defined media for a second overnight growth to synchronise growth. Cells were then diluted in fresh media and, after 4- to 6-hour growth, imaged by microscopy. In case of experiments involving different growth media compositions, cultures were inoculated once more at a 100-fold dilution in the appropriate AB media composition for overnight growth and a final re-inoculation in fresh media with dilutions ranging from 2-fold to 100-fold, depending on the overnight attained optical densities. Note that cultures grown under different nutrient conditions attained different densities after overnight growth. Care was therefore taken to dilute each culture in its appropriate media such as to obtain adequate numbers of cells for experimental assays but at the same time avoiding saturation of the faster growing cultures. After a final 4- to 9-hour growth period cells were either imaged with fluorescence microscopy or subjected to competition assays.

### Cloning of constructs for genomic insertions and deletions

All primers, plasmids and strains used for cloning and PCR verifications together with brief description of the constructions are listed in Supplemental Table S1. Genomic deletions and insertions in the *X. citri* genome were all constructed using a two-step allelic exchange procedure (Hmelo et al. 2015). For this, 500 to 1000 base pair-sized fragments up- and downstream from the region of interest were amplified using a high-fidelity polymerase (Phusion, Thermo Scientific) and cloned into the pNPTS138 vector either by traditional restriction digest cloning (NEB and Thermo Scientific) or by Gibson assembly (NEB). The resulting plasmid was used to transform the appropriate *X. citri* strain by electroporation (2.0 kV, 200 Ω, 25 µF, 0.2 cm cuvettes; Bio-Rad)(Sawitzke et al. 2011). A first recombination event was selected for on LB plates containing 50 µg/ml kanamycin. Transformants were streaked for single colonies on kanamycin plates whereafter several single colonies of the merodiploids (Kan^R^, Suc^S^) were streaked on sucrose plates selecting for a second recombination event creating either a wild-type or mutant allele. After confirmation of the loss of the kanamycin resistance cassette together with *sacB*, a PCR was performed using primers that hybridize outside of the homology regions to identify the target allele. Strains containing the wild type alleles (revertants created at an equal rate during the second recombination event) were also stored and used as controls for their respective mutants. For the insertion of the pPM7G plasmid into the *amy* gene of *X. citri,* cells were electroporated with the pPM7G plasmid and selected for on LB plates containing 50 µg/ml kanamycin. Integrity of the *virB* operon in these strains was confirmed by PCR to exclude erroneous recombination with the pPM7G cloned 5’UTR_B7_ regions.

### Chlorophenol red-β-D-galactopyranoside (CPRG) bacterial competition assay

To visualize and quantify the ability of *X. citri* to lyse *E. coli* strain MG1655, a CPRG-based method was used as described previously (Vettiger & Basler 2016; Sgro et al. 2018). Briefly, to each well of a clear U-shaped bottom 96-well plate, 100 µL of a mixture of 0.5 X buffer A (7.5mM (NH_4_)_2_SO_4_, 8.5mM Na_2_HPO_4_, 11mM KH_2_PO_4_, 25mM NaCl, pH 7.0), 1.5% agarose and 40 µg/mL CPRG (Sigma-Aldrich) was added, and plates were thoroughly dried under a laminar flow. *X. citri* cells grown in the appropriate media, were mixed in a 1:1 volume ratio with a concentrated *E. coli* culture. The *E. coli* cultures were grown to OD_600_ = 1 in the presence of 0.2 mM IPTG (inducing the *lac* operon) in 2xYT medium, washed once and concentrated 10 times. Five microliters of *X. citri* and *E. coli* mixtures were immediately added to the 96-well plate without puncturing or damaging the agarose, covered with a transparent seal and quickly thereafter absorbance at 572 nm (A_572_) was monitored over time in a 96-well plate reader for at least 200 minutes (SpectraMax Paradigm, Molecular Devices). The A_572_ values were processed using RStudio software (RStudio-Team 2016) and plotted using the ggplot2 package (Wickham 2016). Background intensities obtained from the mean of A_572_ values of non lysing *E. coli* cells were subtracted from the data series and data were normalized for initial OD_600_ differences.

### Fluorescence microscopy image acquisition and analysis

Briefly, 1 µL of cell suspension was spotted on a thin agarose slab containing 1X buffer A (15mM (NH_4_)_2_SO_4_, 17mM Na_2_HPO_4_, 22mM KH_2_PO_4_, 50mM NaCl, pH 7) and 2% agarose and covered with a #1.5 cover glass (Corning). For time-lapse imaging, thicker agar slabs containing the appropriate media were constructed as described (Bayer-Santos et al. 2019). Phase contrast and msfGFP emission images were obtained with a Leica DMI-8 epifluorescent microscope. msfGFP emissions were captured using 1000 to 1500 ms exposure times at maximum excitation light intensities. The microscope was equipped with a DFC365 FX camera (Leica), a HC PL APO 100x/1.4 Oil ph3 objective (Leica) and a GFP excitation-emission band-pass filter cube (Ex.: 470/40, DC: 495, Em.: 525/50; Leica). For detection of VirB10-msfGFP foci, eleven 0.05 µm Z-plane stacks were obtained from a 0.5 µm region within the centre of the cells. This allowed for a better signal to noise ratio of the VirB10-msfGFP foci and increased detection of VirB10-msfGFP foci location in different depths of the cell. These image stacks were background subtracted by a rolling ball correction using a significant cell-free portion of each image as a reference and, finally, combined by an average intensity projection using the FIJI software package (Schindelin et al. 2012). To obtain a quantitative representation of cell sizes, background corrected fluorescence intensities and amount of foci present per cell, the images were analysed using the MicrobeJ software package (Ducret et al. 2016) and data was analysed by RStudio software (RStudio-Team 2016) and plotted using the ggplot2 package (Wickham 2016).

### Co-culture growth experiment

To illustrate the physiological burden associated with T4SS overexpression seven independent *X. citri ∆5’UTR*_*B7*_ *virB11-msfgp* mutants and seven independent *X. citri virB11-msfgfp* strains (revertants to wild-type from the second recombination event), in two separate experiments, were grown overnight in 2xYT media, diluted 100-fold into defined AB media with 0.2% sucrose and 0.2% casamino acids and grown overnight. These overnight cultures were mixed 1:1 and 10-fold diluted into fresh AB media with 0.2% sucrose and 0.01% casamino acids. Immediately after mixing (at timepoint 0h) fluorescence microscopy images were taken (as described above) to register the exact ratio of *X. citri virB11-msfgfp* cells versus *X. citri ∆5’UTR*_*B7*_ *virB11-msfgfp* cells. The cultures were subsequently diluted regularly so to prevent cultures from reaching saturation which would halt further cell division. At the indicated time-points several microscopy images of each of the co-cultures were again acquired. Given that the *virB11-msfgfp* reporter in the strain lacking the *5’UTR*_*B7*_ has a higher msfGFP production (histograms of the populations’ msfGFP fluorescence levels do not overlap), cells could be sorted by using average msfGFP content and as such, an accurate quantification of the cell ratio between wild-type and deleted *5’UTR*_*B7*_ strains could be calculated over time.

### Transcription start site analysis

Transcriptional start sites of the *virD* and *virB* transcripts were obtained by using the 5’ RACE Kit (Roche), as described (Andrade et al. 2014). The oligos for *virD4* and *virB7* 5’ RACE assays are listed in Supplemental Table S1. The resulting PCR fragments were blunt ligated into pGEM-T before sequencing 3 independent clones, identifying the transcription start sites.

### RNA electromobility shift assays

For the RNA electromobility shift assays, DNA fragments encoding either the entire 5’ UTR of the *virB* operon or a shortened fragment lacking the first 73 nucleotides (∆1-73nt), were amplified from *X. citri* genome using forward primers which include the T7 promoter sequence (see Supplemental Table S1). RNA transcripts of the cloned *5’UTR*_*B7*_ fragments were produced *in vitro* from the resulting purified PCR products by using the T7 Transcription kit (Roche) and labeled by using the RNA 3’ end biotinylation kit (Pierce). The CsrA recombinant protein was purified as previously described (Andrade et al. 2014). Approximately 70 nM of purified CsrA protein and 6.25 nM Biotin-labeled RNA were mixed with binding buffer [(10 mM HEPES pH 7.3, 20 mM KCl, 1mM MgCl2, 1 mM DTT, 5% glycerol, 0.1 µg/µL yeast tRNA, 20 U RNasin (Promega)] in a total reaction volume of 20 µL. The binding reactions were incubated at 25°C for 20 min. A 5 µL aliquot of loading buffer (97% glycerol, 0.01% bromophenol blue, 0.01% xylene cyanol) was added to the binding reaction and immediately loaded and resolved by 5% native polyacrylamide gels. The binding assays and detection of RNA products were performed with the LightShift Chemiluminescent RNA EMSA Kit (Thermo Scientific). For the control reactions, 312.5 nM competitor unlabeled RNA of the *virB 5’UTR*_*B7*_ was added to the binding reactions.

## Acknowledgments

We thank Alexandre Bruni-Cardoso for unlimited access to the Leica DMI8 microscope. Special thanks to Ioannis Passaris and Sander K. Govers for very helpful discussions, suggestions and critical reading of the manuscript, and to Gilberto Kaihami for the introduction to the R programming language. This work was supported by a FAPESP post-doctoral grant to William Cenens (2015/18237-2) and FAPESP research grants to Chuck S. Farah (2011/07777-5 and 2017/17303-7).

## Supplementary File Legends

**Supplemental Table S1**: Primers, strains and plasmids used in this study

**Supplemental Figure S1**: 5’RACE assay identifying the transcription start sites for *virD4* and *virB7*. **a)** bottom panel shows the two adjacent possible transcription start sites of *virD4* (red and purple arrow) due to sequencing ambiguity. The possible start sites are indicated with a purple G and a red C in the nucleotide sequence (top panel). *VirD4* sequence starts at the end of the displayed nucleotide sequence. In between the TSS and *virD4* a putative ORF of unknown function is present. **b)** Similar analysis for *virB7* with start site depicted with red arrow (bottom panel) and red G in the nucleotide sequence (top panel). In panel a and b, translation start sites are depicted in red and open reading frames in blue. In bold are the putative polymerase binding sites and in underlined regular font the ribosome binding sites.

**Supplemental Movie S1**: Time-lapse movie showing contact dependent lysis at the single-cell level during growth in media depleted for casamino acids. Movie starts after 25 hours of growth on the pad. Clearly showing greatly decreased growth speeds. White arrows indicate regions were *E. coli* cells come into contact with the smaller sized *X. citri* cells. Scalebar indicates 5µm. Timestamp in bottom left of the movie.

**Supplemental Movie S2**: Time-lapse movie showing contact dependent lysis at the single-cell level during growth in rich media. Movie was started and ran in parallel with Movie S1. Starting at timepoint zero. White arrows indicate regions were *E. coli* cells come into contact with smaller sized *X. citri* cells. Scalebar indicates 5µm. Timestamp in bottom left of the movie.

## Notes

#### Summary of Updates

Author affiliations updated, small typos corrected

