## Supplemental Figure S1 for "Bactericidal Type IV Secretion System Homeostasis in Xanthomonas citri"

**a**

### Transcription start site of *virD4*

GGTCAGCGCTGCCTACCGCAGCCCAGTCCGCCGATACC**GCTTTC**ACCGTCACACGTCTTGCT
T**TAGCAT**CGCCACC**GC**GCTGACGGAAAGGAAGGCTGGCG**ATG**GACGAATATCAGCACACTGT
GCTGACCAGGGGGGGTACCGTGTTGTGGCGATTACGCGTGACGAGGTGTACGCACCCGACG
CCGTCGTGGCATACGCCGTTGTGACCGACGCGGGTACGCAATCACTCCGGATTTGTCCCTC
GACCAGGCCAAGGTCTGGATCGATTCGCTGGTGGAGAGCGAGAGTGGCGGGCGCAAGTCCGG
CCTTGTCGACCATAAGCCGGTCGTGCGTCGCTAGCCCGCCATCGTGATGCAAGCCTAAGGAG
AAGGGCGC**ATG**TCCAAACCCAAAATCATCGCAATTGCCACGCTGGTGG

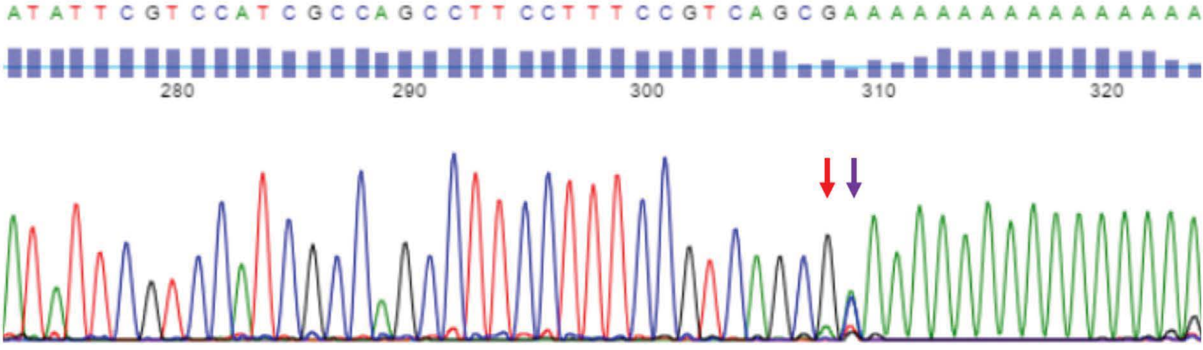


**b**

### Transcription start site of *virB7*

TGTCTCGTTTGCAGCGCAGGGCTTAGCGCGCAATT**CTTGAC**GGATCTGCTTGTTAAGGGC**AAAGTT**
AGCAA**G**AGTCTGCGGTTTAGATCGATGGTCGTAGGGGAGCGGCGCCGGATGGGGCCGTACTGGAGT
AAGGCTATGGATGGTTTGTCAGCGTATTCATTTTTTCCAGCAGCACACACTTGCAACTGGGGCGAC
CCAGCGGGTCGTAGGACTGCGAAAAACGCATCAATAACAAGTTTCCTCCGGTACCCACTGGTGCGC
GGAGGCTTAGTGCTGTCCGTCTTAAGCCCGAAATCTCATGAGTTGGAGAAGTCAGA**GTG**AATCCGA
TGTATGTGTCCAAGCTGTCGTTGGTGCTAGTGGCTG

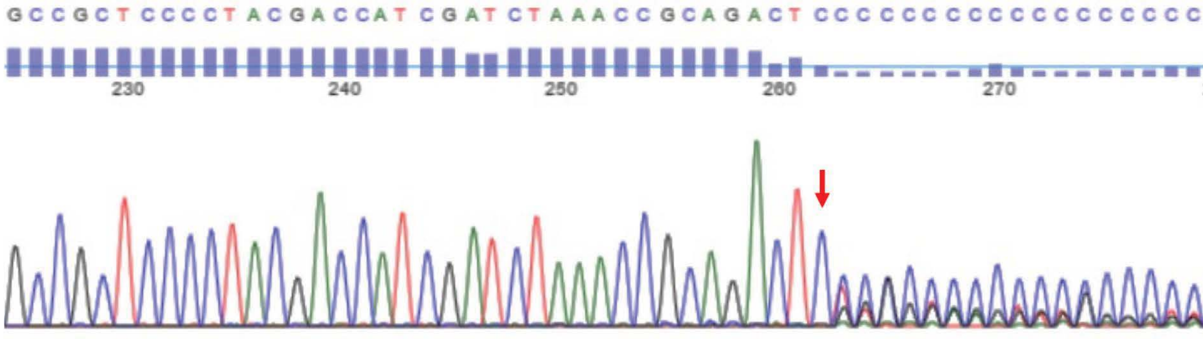
