## Supplementary figures and images for "Bactericidal Type IV Secretion System Homeostasis in Xanthomonas citri"

### Supplemental Movie S1

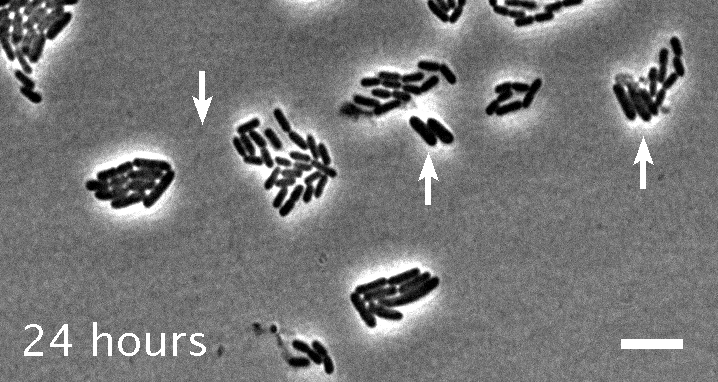

### Supplemental Movie S2

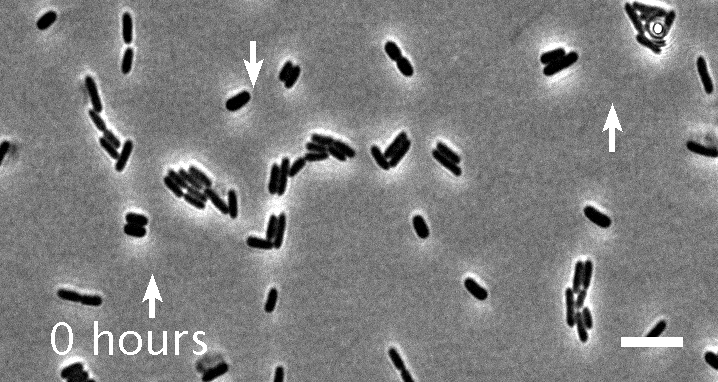
